# Coexpression reveals conserved mechanisms of transcriptional cell identity

**DOI:** 10.1101/2020.11.10.375758

**Authors:** Megan Crow, Hamsini Suresh, John Lee, Jesse Gillis

## Abstract

What makes a mouse a mouse, and not a hamster? The answer lies in the genome, and more specifically, in differences in gene regulation between the two organisms: where and when each gene is expressed. To quantify differences, a typical study will either compare functional genomics data from homologous tissues, limiting the approach to closely related species; or compare gene repertoires, limiting the resolution of the analysis to gross correlations between phenotypes and gene family size. As an alternative, gene coexpression networks provide a basis for studying the evolution of gene regulation without these constraints. By incorporating data from hundreds of independent experiments, meta-analytic coexpression networks reflect the convergent output of species-specific transcriptional regulation.

In this work, we develop a measure of regulatory evolution based on gene coexpression. Comparing data from 14 species, we quantify the conservation of coexpression patterns 1) as a function of evolutionary time, 2) across orthology prediction algorithms, and 3) with reference to cell- and tissue-specificity. Strikingly, we uncover deeply conserved patterns of gradient-like expression across cell types from both the animal and plant kingdoms. These results suggest that ancient genes contribute to transcriptional cell identity through mechanisms that are independent of duplication and divergence.

## INTRODUCTION

Understanding how the genome changes as species diverge is a central question in evolution. With access to sequence data, comparative genomics research initially focused on the association between gene family conservation and the phenotypes that emerge in particular lineages^1–4^. While this approach continues to shed light on genome evolution^5^, it provides at best an incomplete picture, omitting phenotypic differences that can be driven by changes in gene regulation^6,7^. There has been growing interest in using functional genomics data to find regulatory differences by comparing homologous samples, often focusing on gene expression as the output of changing regulatory architecture between species^8–12^. Yet because of its dependence on anatomical homology, this approach is necessarily limited to more closely related species. How then, can we compare regulatory conservation across the tree of life? The answer is coexpression.

In a coexpression network, genes are nodes and the edges represent expression similarity between genes, often a correlation coefficient. Functionally related genes are often adjacent in the network as their expression is coordinated across biological conditions, allowing for the inference of gene regulatory modules through clustering. As early as 2003, comparative coexpression approaches were used to demonstrate similarity in gene regulation between species as distant as plants and mammals^13–18^. However, these studies were limited by both the data and methods available, often relying on a small number of microarray datasets from a handful of model organisms, and ortholog predictions from BLAST^19,20^.

We breathe new life into this area by taking advantage of high-powered coexpression networks from animals, plants and yeast RNA-sequencing (RNA-seq) data^21^, as well as modern orthology prediction algorithms^22–24^, and measuring the conservation of coexpression relationships. As expected, we find that coexpression conservation tracks with evolutionary distances^25^, and is significantly higher for genes expressed in all cell types. However, by linking these results to single-cell expression data^26–30^, we find a fascinating possible role for “ubiquitous” genes in transcriptional cell identity. In both mouse and *Arabidopsis,* we find constitutively expressed genes with gradient-like expression distributions across cell types. In combination with their high degree of co-expression conservation, this suggests that these genes have tightly maintained functions that contribute to continuous aspects of cell identity. Coexpression conservation provides a data-driven estimate of gene functional divergence across species, and we have made all of our data, methods and results available to facilitate its use.

## RESULTS

### Establishing meta-analytic coexpression networks as a tool for comparative genomics

Reliable estimates of species-specific gene coexpression patterns are a necessary backbone for comparative analysis of regulatory divergence. We recently published a set of networks with our coexpression webserver, CoCoCoNet^21^, which contain RNA-seq data from 14 species across 895 datasets, and more than thirty-nine thousand individual samples (**Figure 1A**). Here, we establish the power and robustness of these networks by measuring the connectivity of genes with the same Gene Ontology (GO) annotations^31^, and by evaluating the stability of results after bootstrapping the network building process.

**Figure 1.**
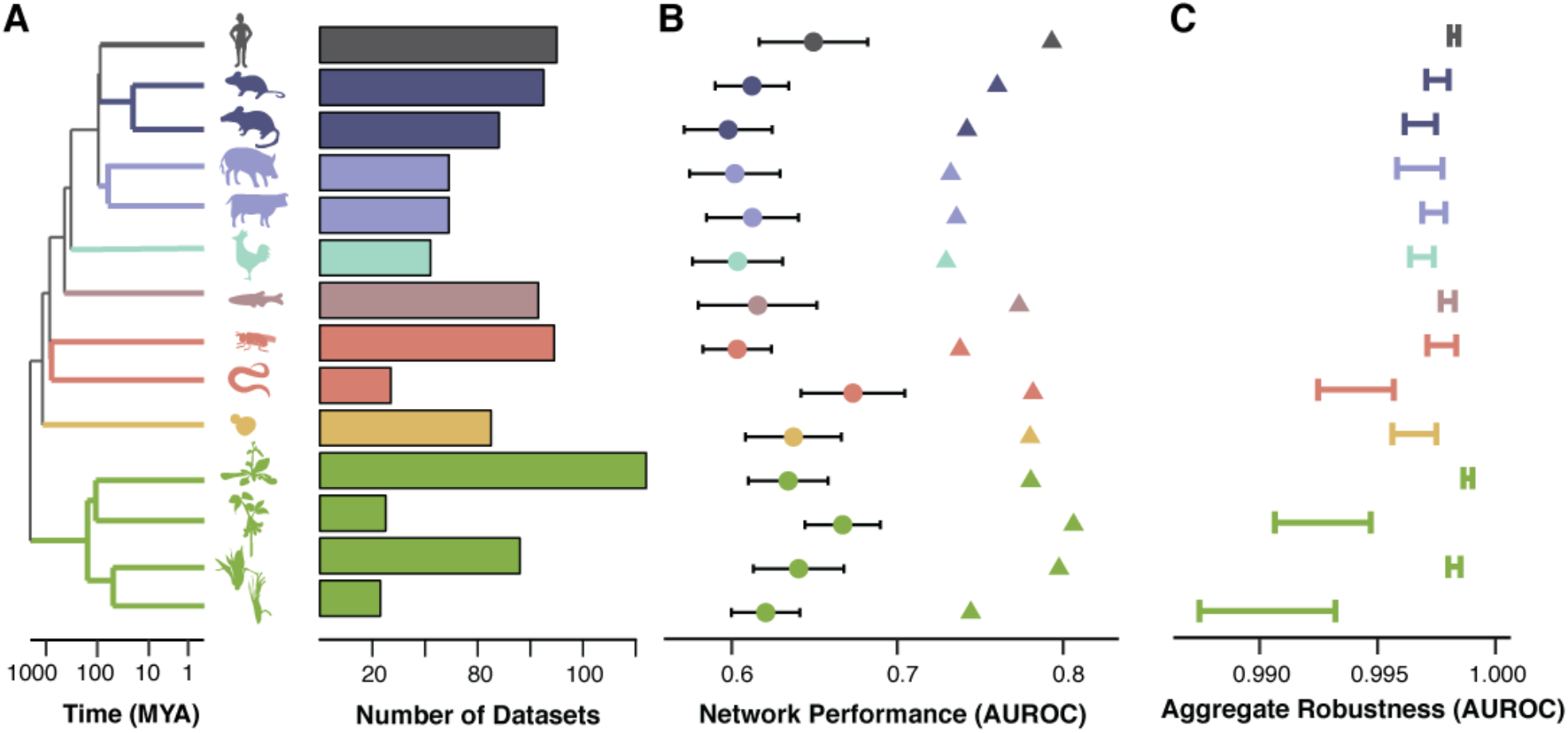
Aggregate coexpression networks are a powerful tool for comparative genomics. **A** – CoCoCoNet contains data from 14 species in three kingdoms: plants, animals and fungi. The dendrogram shows phylogenetic relationships between these species, and the barplots indicate the number of datasets used to build aggregate coexpression networks. **B** – Circles show the mean GO prediction performance for individual networks (+/− standard deviation) while triangles indicate aggregate network performance. **C** – Aggregate robustness is high across all species, with variation dependent on n.

As a first validation, we find that species-specific networks built from multiple datasets (“aggregate” networks) have strong connections between genes from the same GO group, and that these connections are significantly stronger than those found in networks from individual datasets (**Figure 1B**, neighbor voting mean Area Under the Receiver-Operating characteristic Curve (AUROC) individual networks=0.63, mean AUROC aggregates=0.76, Wilcoxon p<10^−8^, n=14 species). To evaluate the robustness of the aggregate networks, we bootstrapped the aggregation procedure 10 times, then used the ranked edges in the bootstrapped networks to predict the top 1% of edges in the reference aggregate networks (**Figure 1C**). Performance at this task was close to perfect (mean AUROC = 0.996 +/− 0.002), while variability between bootstrapped and reference aggregate networks declined as a function of the number of experiments and samples as expected (Spearman correlation coefficient=−0.83 for experiments, −0.86 for samples). These results indicate that aggregate networks are statistically robust and biologically meaningful, with strong connections between functionally related genes.

### Similarity of coexpression neighborhoods quantifies ortholog conservation

Having established the robustness of our networks, we next explore the degree to which they can be used for cross-species comparisons. Our analysis focuses on characterizing the degree to which pairs of orthologs have retained similar coexpression patterns, or “neighborhoods” in the networks.

To measure the similarity of ortholog neighborhoods between two species, we first subset networks to include only one-to-one orthologs between that species pair (e.g., pig and yeast as shown in the schematic, **Figure 2A**). Next, each gene’s neighborhood is defined by ranking all edges associated with it, and then the ranks of the gene’s top coexpressed gene pairs are compared across species (see Methods for details). This is expressed as an AUROC and so it ranges from 0-1, with 1 meaning perfect coexpression conservation, 0.5 consistent with random re-ordering of neighborhoods, and 0 meaning that coexpression partners have inverted from being the top ranked to bottom ranked across species.

**Figure 2.**
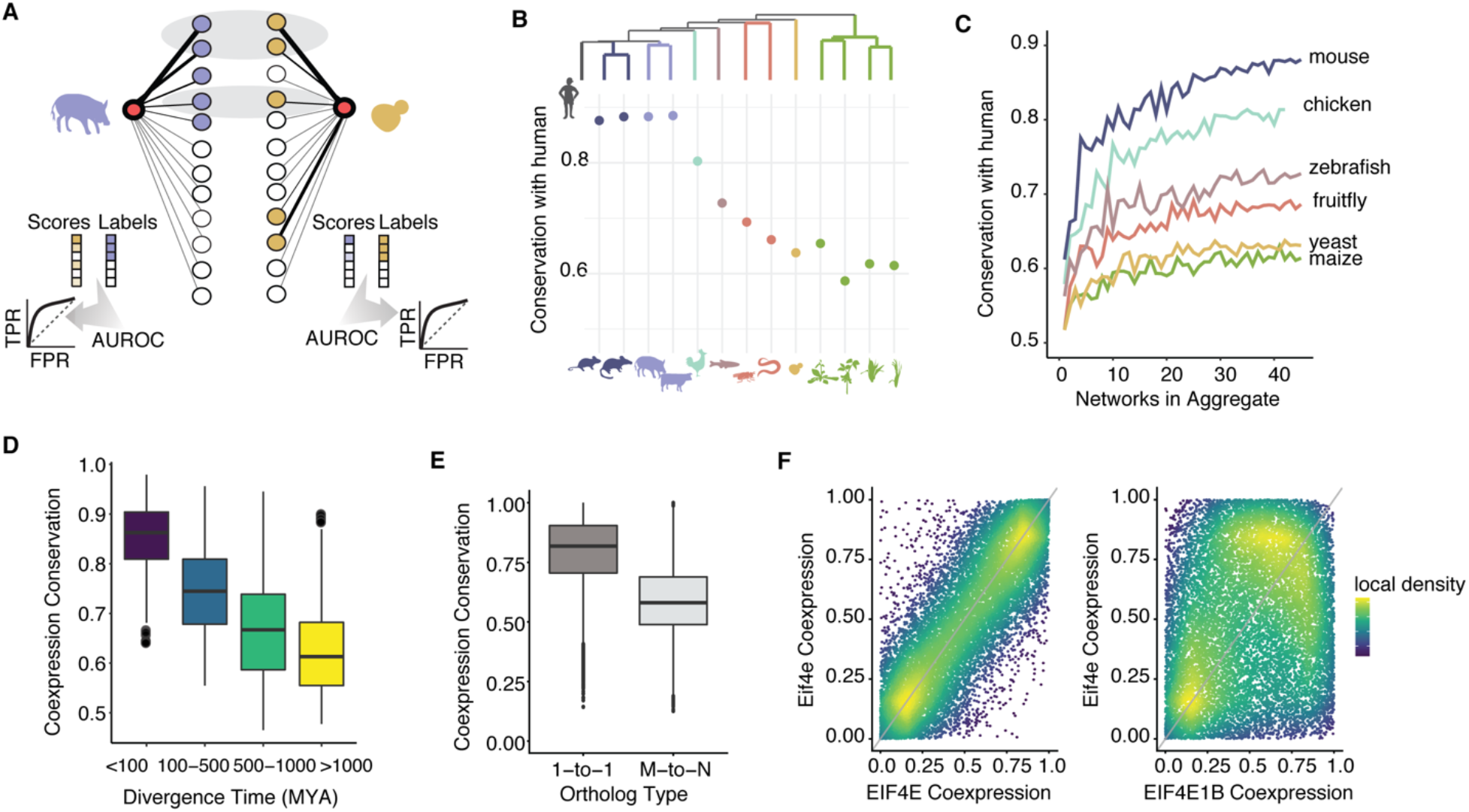
Coexpression divergence correlates with phylogeny and orthogroup size. **A -** Method schematic. Circles represent genes and line thickness indicates strength of coexpression between one target gene (red) and all others. For each target gene in pig, we identify the set of pig genes that are maximally coexpressed with it, shown in blue. We evaluate how conserved this coexpression pattern is in yeast, then repeat the task in the other direction. Genes with high coexpression to the target in both species are highlighted in the gray ovals. Coexpression conservation is reported as the average AUROC in both directions (i.e., pig-yeast and yeast-pig). **B -** Points show mean coexpression conservation for 1-to-1 orthologs between human and each other species. Coexpression conservation is negatively correlated with phylogenetic distance (rho=-0.95, p<10^-6^). **C -** Mean coexpression conservation for 1-to-1 orthologs between human and each of the listed species are plotted against the number of networks included in the aggregate network. Performance increases with additional data. **D -** Boxplots show coexpression conservation scores for 492 orthologous groups defined at the last common ancestor of all eukaryotes, plotted with respect to species divergence times. As in panel B, coexpression is more conserved among more recently diverged species. **E** – Boxplots show coexpression conservation scores. 1-to-1 orthologs are more conserved than N-to-M orthologs (Wilcoxon p<10^−16^). **F** – Coexpression profiles for a 1-to-2 mouse-human ortholog group. The mouse gene Eif4e has a strongly conserved coexpression profile with human EIF4E (left, coexpression conservation AUROC=0.94) but not with human EIF4E1B (right, AUROC=0.41).

As a first validation of our approach, we find that coexpression conservation is strongly negatively associated with phylogenetic distances between species, demonstrated in **Figure 2B** with respect to distance from human (Spearman correlation coefficient= −0.95). We also find that coexpression conservation is sensitive to the amount of underlying data as expected, with strong performance achieved with the inclusion of twenty datasets and scores plateauing beyond this point (**Figure 2C**, mean individual networks=0.55+/−0.04, mean 20-dataset aggregate=0.68+/−0.08, Wilcoxon p<0.002, n=7 species).

One-to-one orthologs are frequently used for comparative genomics analyses^32^, however, as species grow more distant to one another, there are fewer one-to-one orthologs for comparison, particularly in the plant kingdom, where genome duplication events are common^33^. To explore more distant and complex relationships, we generalized our method to be able to compare groups of orthologs, i.e., all genes descended from a single gene in a common ancestor, including lineage-specific duplicates (see **Supplementary Figure 1** for a schematic). As validation, we assessed the conservation of groups of orthologs descended from the last common ancestor of all eukaryotes. We find that coexpression conservation scores for many-to-many (aka N-to-M) orthologs are strongly associated with phylogenetic distances between species (**Figure 2D**). We next evaluated all N-to-M orthologs defined in the last common ancestor between each pair of species, finding that one-to-one orthologs have higher conservation than N-to-M orthologs on average (**Figure 2E**, mean 1-to-1 = 0.79 +/− 0.14, mean N-to-M = 0.59 +/− 0.14, Wilcoxon p<10^−16^). Notably, we also find cases where N-to-M orthologs are strongly differentially conserved, with ~7% of all N-to-M groups containing orthologs with scores >0.7 and <0.5. An example of this is shown in **Figure 2F**. Here, we see that mouse *Eif4e* shares a large fraction of its coexpression neighborhood with human *EIF4E*, but that it is quite distinct from human *EIF4E1B*.

These results indicate that coexpression neighborhoods can be usefully compared across species to provide a measure of ortholog conservation. Distinguishing orthologs by differential coexpression conservation may provide a route forward for discovering genes with “the same” function across species (a.k.a. “functional analogs”).

### Genes expressed in all cell types have conserved coexpression relationships and contribute to cell identity

In the previous section we established that coexpression conservation tracks with phylogeny as expected, and that one-to-one orthologs are more likely to have similar coexpression than those that have duplicated. We also find evidence of strong divergence within orthogroups, which has long been postulated to be an evolutionary mechanism for morphological innovation^34^. Earlier work to probe this hypothesis showed that older genes tend to be more ubiquitously expressed across cell types and tissues, and have greater conservation across species^9,12^, and this has typically been interpreted to mean that younger genes are required for phenotypic novelty.

By leveraging coexpression relationships we can 1) extend these observations to any species pair of interest without requiring knowledge of homologous tissues or cell types, and 2) identify conserved relationships between genes, reflecting conserved regulation and function. Importantly, we can investigate the role of conserved coexpression in cell phenotypes by taking advantage of single-cell RNA-seq data. We use comprehensive single-cell RNA-seq data from the Tabula Muris project for mouse^26^, and four single-cell RNA-seq datasets from *A. thaliana* root^27–30^ as references for cell-type specific expression (**Figure 3A**, see Methods for details), and we use data from the Genotype-Tissue Expression Project (GTEx)^35^ as a reference for human tissue-specific expression.

**Figure 3.**
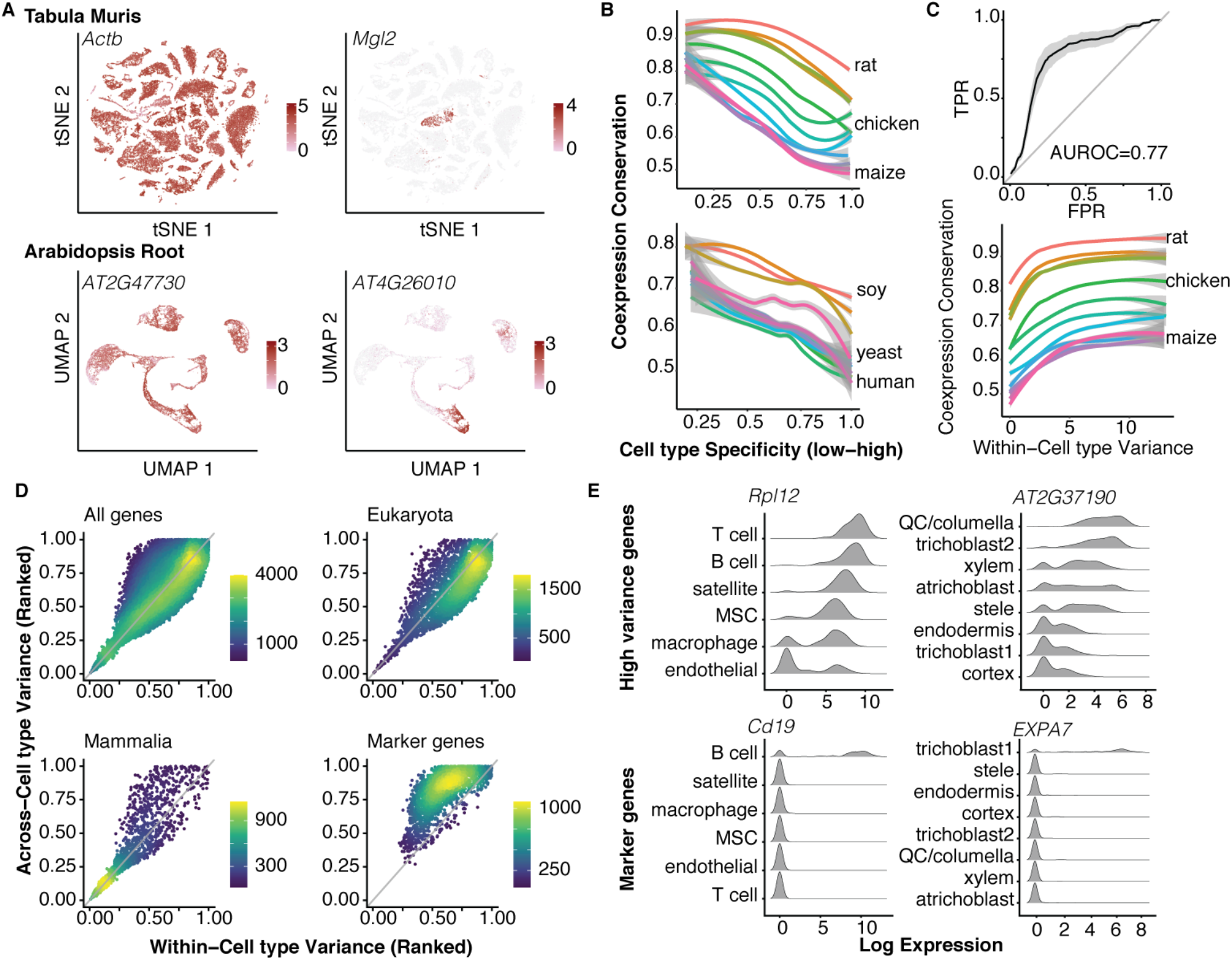
Coexpression conservation suggests ancient, continuous axes of cell identity. **A –** Plots of mouse and Arabidopsis scRNA-seq data, with examples of constitutive (left) vs. cell-type specific expression (right), color indicates expression level. **B –** Coexpression conservation is plotted with respect to cell type specificity for mouse (top) and Arabidopsis (bottom). Lines are loess fits on mean values for each species, +/− SD. Cell type specificity is negatively associated with coexpression conservation. **C –** Expression variance is associated with coexpression conservation. (Top) Expression variance across >200 yeast datasets predicts coexpression conservation. TPR = true positive rate, FPR = false positive rate. ROC curves for all species were binned along x-axis, mean +/− SD is plotted. (Bottom) Coexpression conservation is plotted with respect to within-cell type variance in mouse, with loess fits on mean values for each species. **D –** Across vs. within-cell type variance in mouse is plotted. Colors indicate local point density. The space can be broken into three regions: genes with high-within and high-across variance are typically more ancient, those that are low/low are more recent, and markers have high-across and low-within cell type variance. Due to their high coexpression conservation, high/high genes may have conserved functions vis-à-vis cell identity. **E -** Examples of continuous (top) vs. marker-like (bottom) expression in mouse (left) and Arabidopsis (right). MSC = mesenchymal stem cell, QC = quiescent center. The conserved gene with high within- and across-cell type variance is more continuous across cell types, whereas the markers (high/low) are either on or off.

Consistent with previous research, we find that genes expressed ubiquitously across tissues have higher coexpression conservation than those with tissue-specific expression (**Supplementary Figure 2**, Spearman correlation coefficient = −0.41 +/− 0.06). Expanding this analysis to cell types, we again see the same pattern, with cell-type specificity associated with decreased coexpression conservation in both *A. thaliana* and mouse (**Figure 3B**, mouse Spearman correlation coefficient = −0.50 +/− 0.04, Arabidopsis = −0.26 +/− 0.07). We note that this trend holds across all species pairs used to calculate coexpression conservation. We also confirm that cell type-specific expression and coexpression conservation are associated with estimates of gene age in both kingdoms (**Supplementary Figure 2,** mouse cell-type specificity vs. gene age^36^ Spearman correlation coefficient = 0.47, coexpression conservation vs. gene age = −0.15 +/− 0.1 across 13 species pairs, Arabidopsis cell-type specificity vs. gene age^37–40^ Spearman correlation = 0.27 +/− 0.1, coexpression conservation vs. gene age = −0.23 +/− 0.05, averaged across 6 gene age estimates and 13 species pairs) and that coexpression conservation is significantly higher for orthologs of genes known to be essential in human^41^ (**Supplementary Figure 2,** essential = 0.83 +/− 0.1, non-essential = 0.69 +/− 0.08, Wilcoxon p<0.01, n=13 species compared to human). In summary, gene duplication events are associated with divergence in cell phenotype in both kingdoms, and this is reflected in the divergence of gene-gene relationships among younger and more cell type-specific genes.

But what drives variability in coexpression conservation in species without cell types? Our compendium of networks includes one single-celled organism, the budding yeast *Saccaromyces cerevisiae*. To explore how these results might generalize, we performed a meta-analysis of yeast microarray expression data^42^, using bulk expression variance as an analog for expression in all cell types since they are strongly correlated (**Supplementary Figure 2**). Consistent with our findings in plants and animals, we find that expression variability in yeast is predictive of coexpression conservation (**Figure 3C**, mean AUROC = 0.77 +/− 0.04 SD).

Because variability occurs in the absence of cell type variation in yeast (it may reflect temporal or state variation), we wondered whether we could find a similar effect in multicellular organisms when holding cell type constant. Our expectation is that cells of the same type may vary by cell cycle phase, nascency, or activation state, for example. Strikingly, after calculating expression variation within each cell type in Tabula Muris, we find that the most variable genes have the most strongly conserved coexpression patterns (**Figure 3C**). In other words, while their coexpression partners remain consistent, their expression levels vary dramatically within and across cells. These properties suggest that these genes could contribute to continuous aspects of cell identity that require tightly coordinated signaling, like differences in size or metabolic activity. Cells of different types would likely have broad but distinct expression distributions, forming a gradient of expression when viewed across types. Such “continuous axes” of variation would be more likely to generalize across kingdoms than poorly conserved marker genes (**Figure 3D**).

One example of a gene that might contribute to continuous aspects of cell identity is *Rpl12*, a component of the large ribosomal subunit 60S in mouse. This gene is among the most variably expressed genes within and across cell types (mean standardized rank within = 0.83, across = 0.97) but its expression distribution across cell types is continuous (**Figure 3E**). Given *Rpl12*’s molecular function, we speculate that protein synthesis rate could be a continuous axis of variation from which all cells sample. Notably, we find a similar pattern of continuous expression when we look at an Arabidopsis ortholog of *Rpl12* (**Figure 3E**). This pattern is in stark contrast to more typical marker genes, which tend to have high variance across cell types, and lower variance within cell types because they are “switch-like” in their expression patterns. Two examples are shown in **Figure 3E**: *Cd19*, which is exclusively expressed by mouse B-cells and is conserved only among mammals (mean standarized rank within = 0.55, across = 0.97); and *EXPA7*, which is exclusive to Arabidopsis root trichoblasts and is conserved only among eudicots (mean standardized rank within = 0.77, across = 0.99).

In summary, our combined analysis of single-cell RNA-seq and yeast expression variation shows that while cell type- or tissue-specific marker genes are generally not well conserved, genes with high expression variability are deeply conserved, and may contribute to continuous aspects of cell transcriptional identity, in both single-celled and multicellular organisms.

### Coexpression conservation is associated with ortholog confidence and can predict functional analogs

Our coexpression conservation analysis is made possible by the use of OrthoDB to define orthologous genes. However, orthology prediction is an active area of research in genomics^23,43^, and relying on a single algorithm has known limitations^44–46^. How would our results hold up if we had used a different reference? In the following, we take advantage of orthology information from the Alliance of Genome Resources^24^ which has predictions from 12 independent sources^47–58^ for humans and 6 common model organisms. We assess coexpression conservation across algorithms, and we explore how coexpression conservation can be applied to predict functional analogs.

The majority of algorithms within the Alliance of Genome Resources database (9/12) have high concordance across their orthology predictions (**Figure 4A**, mean Jaccard=0.7), with exceptions attributable to differences in species coverage. In keeping with their overall similarity, we find that average scores for these 9 algorithms are close to tied (mean = 0.74 +/− 0.009, **Supplementary Table 1**) and although OrthoDB is fairly distinct in its predictions (mean Jaccard index=0.18), it performs very close to this average (0.73 +/− 0.13). Remarkably, even though the algorithms are tied on average, we find that where they agree, coexpression conservation is preferentially high: for almost all pairs of species, coexpression conservation is correlated with the number of algorithms that predict the relationship (human-worm shown as an example in **Figure 4B**, all correlations in **Figure 4C**). The only exceptions to this rule are among the three mammals, where coexpression conservation is almost uniformly high.

**Figure 4.**
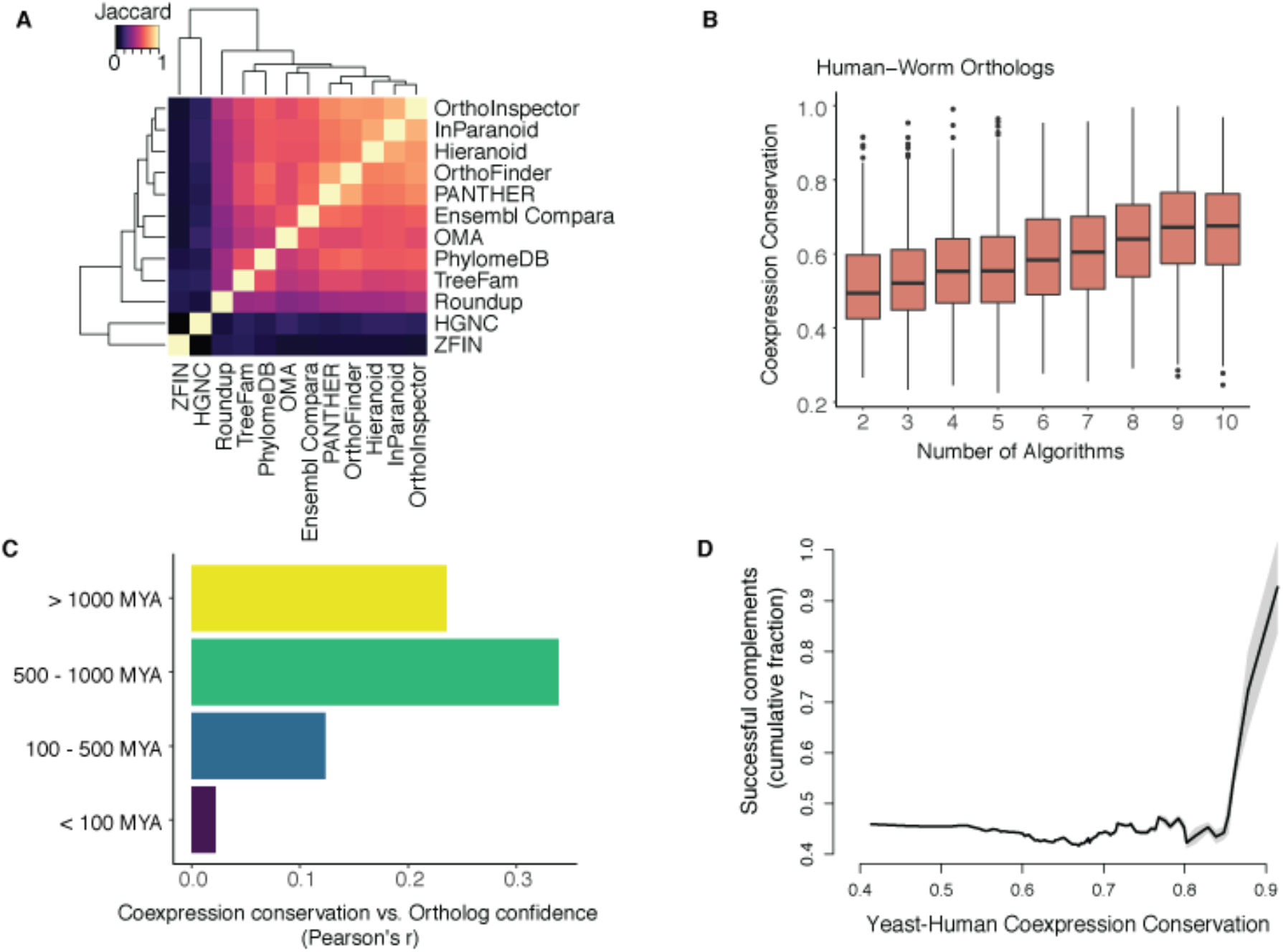
Coexpression conservation is associated with ortholog concordance across algorithms and can predict human-yeast functional analogs. **A –** Heatmap of algorithm concordance. The majority of algorithms (9/12) make similar predictions, with outliers arising from selection biases (i.e., inclusion of only a subset of species). **B -** Mean coexpression conservation for human-worm orthologs is plotted against the number of algorithms predicting the relationship. Coexpression conservation correlates with ortholog confidence. **C –** Bars show the correlation between the number of algorithms and coexpression conservation for each gene pair, binned into three divergence times. Coexpression conservation correlates with ortholog confidence for pairs of species that diverged >100MYA but not for more recently diverged species. **D –** Cumulative success of human-yeast complementation is plotted as a function of coexpression conservation. Ortholog pairs with high coexpression conservation are likely to compensate.

A primary application of orthology prediction is to infer shared function across species, but benchmarks recurrently find a precision-recall trade-off across algorithms^46,59^, with little evidence that any one approach outperforms another^43^. By incorporating functional information directly, coexpression conservation scores may improve sequence-based inference. For example, recent work has found that human genes with similar coexpression patterns can compensate for the loss of their yeast orthologs in complementation screens^60,61^. Here we find that coexpression conservation scores from our independent data and analysis are predictive of this effect (**Figure 4D**). However, we also find that certain pairs of complementing orthologs have very low coexpression conservation scores, with the two lowest scoring pairs excluded from the Alliance of Genome Resources database (**Supplementary Table 2**).

Altogether, these results highlight that confidence in sequence-based orthology is reflected by similarity in gene coexpression relationships, and suggest that a wisdom-of-the-crowds approach may be beneficial for ortholog prediction. Since our earlier results rely only on OrthoDB, they likely represent a lower limit for coexpression conservation which would only improve with the use of meta-orthology methods^62^. Moreover, our approach makes it possible to compare data from any species with sufficient RNA-seq data, not just popular model systems like yeast which have been the focus of many studies due to the greater availability of experimental resources.

## DISCUSSION

By combining sequence-based orthology predictions with robust estimates of gene coexpression neighborhoods, we have developed a measure of gene conservation that can be calculated between any pair of species. To our knowledge our study is the largest of its kind to date, making use of data from hundreds of individual studies, and measuring coexpression conservation across very long diverged species. We find that coexpression conservation is associated with phylogenetic distances, expectations of conservation based on gene family size, and can even predict concordance across orthology prediction algorithms. Moreover, taking advantage of the recent explosion in single-cell data, we identify commonalities between the forces that drive conservation in both single-celled and multi-cellular organisms. The genes that vary most – both within cells of the same type and across cells of different types or states – show deeply conserved patterns of coexpression, suggesting their fundamental role in eukaryotic cell function and identity.

Previous research to investigate the changing gene expression landscape across species focused on the relative lack of conservation among tissue and cell type-specific genes^9,12^, which has been interpreted as evidence for Ohno’s theory that gene duplication and divergence is critical for phenotypic evolution^34^, such as the creation of novel cell types. With our method, we confirm these previous findings, but also extend them, presenting a challenge to Ohno’s theory. Genes with cell type specific expression have more poorly conserved coexpression patterns than those that are expressed in cells of all types, as previously described. Yet genes expressed in all cell types not only have strongly conserved coexpression patterns, but they also have gradient-like expression ****across**** cell types. Variation in activity across cell types is likely what drives their strong coexpression within networks.

What does this mean for cellular evolution and multicellularity? We hypothesize that these genes work in tightly coordinated modules to tune non-negotiable aspects of cellular identity, like cell size, or metabolic rate, generating diversity that allows cells to respond to varying environments. With these diverse populations established, evolution could then use novel genomic variation to mark cell types and refine their organismal roles. Further work to explore this hypothesis is necessary, including additional analyses of single-cell data from long diverged species. However, we note that this will require targeted investigation, as typical single-cell analyses are designed to identify genes that are strongly variable across cell types (i.e., markers) rather than subtler continuous signals. Cell identity is known to be broadly distributed across the transcriptome, and this allows low-depth single-cell RNA-seq to find expected cell clusters even when individual marker genes are not sampled^63–65^. Determining the relative contributions of switch-like versus continuous expression to cell identity will be informative for updating empirical and mechanistic models cell type.

This work is at the intersection of transcriptomics and orthology prediction, and improvements in both will allow for increasing precision of coexpression conservation. Based on our current estimates of available data, we can readily extend our analyses to an additional thirty-eight species. The ability to move beyond that will require efforts to profile the transcriptomes of a greater diversity of organisms, similar to the goals of the Genome10K project for genome sequences^66^. However, a necessary consequence of using bulk RNA-seq data from heterogenous samples is that networks are better powered for genes that are expressed in all cells, and our results make it clear that tissue-specific networks^67^ cannot overcome this as genes expressed in all cell types will continue to dominate. Instead, future work to develop cell-type specific coexpression networks^68^, including methods to map cell identities across distantly related species^69^, and/or to selectively sample the transcriptome for genes with lower expression^70^, could be routes forward. Regarding orthology, one of our most important results is that our conservation measure is correlated with the number of algorithms that predict the orthology relationship. Orthology algorithms are notoriously difficult to benchmark given the lack of gold-standard data (i.e., we lack genomes for the last common ancestors of extant species). Our results suggest that a wisdom-of-the-crowds approach, and incorporation of functional genomics data, could improve on baseline predictions^71^.

Defining the regulatory mechanisms that allow for evolutionary divergence is a central goal in biology. Our method and data now provide a clear route forward for investigating these mechanisms with both breadth and depth.

## METHODS

### Analysis of public gene expression data

All analyses were performed in R. Results are reported as means +/− standard deviations unless otherwise specified.

Aggregate coexpression networks were downloaded from CoCoCoNet^21^. Networks for individual datasets are stored internally and are available on request. In brief, networks for each dataset are built by calculating the Spearman correlation between all pairs of genes, then ranking the correlation coefficients for all gene-gene pairs, with NAs assigned the median rank. Aggregate networks are generated by averaging rank standardized networks from individual datasets. To assess the connectivity of GO groups we used the *run_neighbor_voting* function from the EGAD R package^72^, subsetting GO to terms with 10-1000 genes. For human tissue specificity analyses, processed expression data^73^ from the GTEx project^35^ (mean expression per tissue) was downloaded from github.com/sarbal/EffectSize. All −1 values were assigned NA, then tissue specificity was calculated as published^74^.

The Tabula Muris Smart-seq2 expression matrix and sample metadata were downloaded from <u>FigShare</u> and converted to *SingleCellExperiment* objects for further processing. For each tissue, counts were normalized using the *logNormCounts* function from the scater package^75^. Cell type specificity was calculated as published for each tissue^74^, then cell type specificity scores were averaged across tissues, excluding NAs. To visualize cells, we used the t-SNE coordinates provided by the authors. Within-cell type variance was calculated for each cell type in each tissue separately, then averaged for each tissue, and averaged across tissues. Across-cell type variance was calculated for each tissue separately using the mean log expression for each cell type, then values were averaged across tissues. For Figure 3D, within and across-cell type variance were ranked and standardized between 0-1 such that the highest variance genes would have a score of 1. The limma package^76^ was used to find marker genes with a log fold-change threshold >4 between each cell type and any other within its tissue.

*Arabidopsis thaliana* root single-cell RNA-seq expression matrices and sample metadata were downloaded from the Gene Expression Omnibus^77^ or provided by the authors. Cell type specificity was calculated for each dataset separately, then averaged, excluding NAs. Within-cell type variance was calculated for each cell type in each dataset separately, then averaged across datasets. Across-cell type variance was calculated for each dataset using the mean log expression in each cell type, then averaged across datasets. To visualize cells from all studies, we first used MetaNeighbor^63^ to find replicable clusters, and subset individual datasets to clusters replicating in at least one other study using an AUROC cut-off of 0.7. We used the *multiBatchNorm* function from the batchelor package for initial batch correction^78^. Then, we selected variable genes using the *get_variable_genes* function from MetaNeighbor and used *fMNN* in batchelor on batch corrected data, subset to variable genes. This provided principle components which were used for the UMAP projection of all cells^79^ (20 components were used).

Yeast microarray data were downloaded from the Saccharomyces Genome Database^42^. We included studies with >10 samples. After rank normalizing expression for each sample, we calculated the variance of expression for each gene that was measured in at least 80% of the datasets. Variance was calculated for each dataset separately, then averaged. For the analysis in Figure 3C, we considered all gene pairs with coexpression conservation >0.9 to be true positives and used the ranked average variance to predict these for each species with the *auroc_analytic* function in EGAD^72^.

### Gene annotations and orthology

Gene function annotations were sourced from the Gene Ontology^31^. GO terms and gene associations were obtained by merging data from the NCBI (ftp://ftp.ncbi.nlm.nih.gov/gene/DATA/gene2go.gz) and biomaRt^80^. Associations were propagated based on “is_a” relationships between terms. In addition, essential gene annotations were obtained from the Macarthur lab’s GitHub repository (https://github.com/macarthur-lab/gene_lists), originally sourced from Hart et al^41^. Yeast-human complementation data was downloaded from the supplement of Kachroo et al^60^. Mouse gene age estimates^36^ were downloaded from the Marcotte lab’s GitHub repository (https://github.com/marcottelab/Gene-Ages) and *A. thaliana* gene age estimates were collected from multiple sources^37–40^.

OrthoDB^22^ was used for orthology mapping. For each pair of species, we search for the most recent phylogenetic split, then obtain inferred orthology groups for all genes descended from the common ancestor. These were either filtered to include only one-to-one relationships, or to include all N-to-M orthologous pairs. We also downloaded orthology information from the Alliance of Genome Resources^24^ for assessment with our N-to-M coexpression conservation method, detailed below. We de-duplicated the information so that each gene pair appeared only once, rather than having a direction from a source-target species. Pairs from all algorithms were considered the “universe” of possible orthologs. Species divergence times were sourced from TimeTree^25^.

### Coexpression conservation

For each pair of species to be compared, we filter aggregate coexpression networks to include known orthologous genes, then we compare each gene’s top coexpression partners across species to quantify gene similarity. We treat this as a supervised learning task, using the ranks of the coexpression strengths from one species to predict the top coexpression partners from the second species, and then repeating this task in the opposite direction, finally averaging the scores. We refer to this as a measure of “coexpression conservation” and note that it is formally equivalent to the average area under the receiver operator characteristic curve (AUROC).

To generalize this to the case of N-to-M orthologs, we describe the analysis in greater detail: Consider a gene A1 in species 1 that has two orthologs in species 2 – genes B1 and B2. First, the top ten genes exhibiting the highest coexpression with A1 are chosen. All possible orthologs in species 2 for the set of top 10s of A1 are shortlisted as the “translated top 10s”. Note that since each gene in species 1 can now have one or more orthologs in species 2, the translated top 10s can vary in length. Additionally, some of the top ten coexpressed genes in species 1 may map to the same orthologs in species 2, but we only consider a unique list of orthologs in the translated top 10s for each gene in species 1. This task is repeated in the opposite direction, where the ranks of coexpression strengths of genes in species 2 are used to predict the top coexpression partners of genes from species 1. Scores from both directions are averaged, thereby providing a measure of overall coexpression conservation.

## DATA AVAILABILITY

Data and code to reproduce the coexpression conservation analysis are available through CoCoCoNet (https://milton.cshl.edu/CoCoCoNet/).

## ACKNOWLEDGEMENTS

Funding for the work was provided by the National Institutes of Health (R01 LM012736 and R01MH113005 for H.S., J.L. and J.G.; K99 MH120050 for M.C.). We would like to thank Trygve Bakken, Dave Jackson, Ken Birnbaum and Stephan Fischer for thoughtful feedback on earlier drafts of this manuscript.

## AUTHOR CONTRIBUTIONS

J.G. conceived the project. M.C. and J.G. wrote the paper, designed analyses and interpreted results. M.C., H.S. and J.L. performed computational experiments, with H.S. contributing to algorithmic design, and J.L. contributing webserver integration. All authors read and approved the final manuscript.

## MATERIALS & CORRESPONDENCE

Correspondence and material requests should be addressed to Jesse Gillis.

## REFERENCES

1 Rubin, G. M. et al. Comparative genomics of the eukaryotes. Science (New York, N.Y.) 287, 2204–2215 (2000).

2 Kaul, S. et al. Analysis of the genome sequence of the flowering plant Arabidopsis thaliana. Nature 408, 796–815 (2000).

3 Copley, R. R., Schultz, J., Ponting, C. P. & Bork, P. Protein families in multicellular organisms. Current opinion in structural biology 9, 408–415 (1999).

4 Hutter, H. et al. Conservation and novelty in the evolution of cell adhesion and extracellular matrix genes. Science (New York, N.Y.) 287, 989–994 (2000).

5 Fernández, R. & Gabaldón, T. Gene gain and loss across the metazoan tree of life. Nature Ecology & Evolution 4, 524–533, doi:10.1038/s41559-019-1069-x (2020).

6 King, M. & Wilson, A. Evolution at two levels in humans and chimpanzees. Science (New York, N.Y.) 188, 107–116, doi:10.1126/science.1090005 (1975).

7 Gilad, Y., Oshlack, A., Smyth, G. K., Speed, T. P. & White, K. P. Expression profiling in primates reveals a rapid evolution of human transcription factors. Nature 440, 242–245, doi:10.1038/nature04559 (2006).

8 Su, A. I. et al. Large-scale analysis of the human and mouse transcriptomes. Proceedings of the National Academy of Sciences 99, 4465–4470, doi:10.1073/pnas.012025199 (2002).

9 Brawand, D. et al. The evolution of gene expression levels in mammalian organs. Nature 478, 343–348, doi:10.1038/nature10532 (2011).

10 Patel, R. V., Nahal, H. K., Breit, R. & Provart, N. J. BAR expressolog identification: expression profile similarity ranking of homologous genes in plant species. The Plant journal : for cell and molecular biology 71, 1038–1050, doi:10.1111/j.1365-313X.2012.05055.x (2012).

11 Cardoso-Moreira, M. et al. Gene expression across mammalian organ development. Nature 571, 505–509, doi:10.1038/s41586-019-1338-5 (2019).

12 Alam, T. et al. Comparative transcriptomics of primary cells in vertebrates. Genome research, doi:10.1101/gr.255679.119 (2020).

13 Stuart, J. M., Segal, E., Koller, D. & Kim, S. K. A gene-coexpression network for global discovery of conserved genetic modules. Science (New York, N.Y.) 302, 249–255 (2003).

14 Bergmann, S., Ihmels, J. & Barkai, N. Similarities and Differences in Genome-Wide Expression Data of Six Organisms. PLOS Biology 2, e9, doi:10.1371/journal.pbio.0020009 (2003).

15 Ruprecht, C. et al. Phylogenomic analysis of gene co-expression networks reveals the evolution of functional modules. The Plant Journal 90, 447–465, doi:10.1111/tpj.13502 (2017).

16 Gerstein, M. B. et al. Comparative analysis of the transcriptome across distant species. Nature 512, 445–448, doi:10.1038/nature13424 (2014).

17 Dutilh, B. E., Huynen, M. A. & Snel, B. A global definition of expression context is conserved between orthologs, but does not correlate with sequence conservation. BMC genomics 7, 10, doi:10.1186/1471-2164-7-10 (2006).

18 Chikina, M. D. & Troyanskaya, O. G. Accurate Quantification of Functional Analogy among Close Homologs. PLOS Computational Biology 7, e1001074, doi:10.1371/journal.pcbi.1001074 (2011).

19 Fitch, W. M. Distinguishing homologous from analogous proteins. Systematic zoology 19, 99–113 (1970).

20 Altschul, S. F., Gish, W., Miller, W., Myers, E. W. & Lipman, D. J. Basic local alignment search tool. Journal of molecular biology 215, 403–410, doi:10.1016/s0022-2836(05)80360-2 (1990).

21 Lee, J., Shah, M., Ballouz, S., Crow, M. & Gillis, J. CoCoCoNet: conserved and comparative co-expression across a diverse set of species. Nucleic acids research 48, W566–W571, doi:10.1093/nar/gkaa348 (2020).

22 Kriventseva, E. V. et al. OrthoDB v10: sampling the diversity of animal, plant, fungal, protist, bacterial and viral genomes for evolutionary and functional annotations of orthologs. Nucleic acids research 47, D807–D811, doi:10.1093/nar/gky1053 (2018).

23 Altenhoff, A. M. et al. The Quest for Orthologs benchmark service and consensus calls in 2020. Nucleic acids research 48, W538–W545, doi:10.1093/nar/gkaa308 (2020).

24 Consortium, T. A. o. G. R. Alliance of Genome Resources Portal: unified model organism research platform. Nucleic acids research 48, D650–D658, doi:10.1093/nar/gkz813 (2019).

25 Kumar, S., Stecher, G., Suleski, M. & Hedges, S. B. TimeTree: A Resource for Timelines, Timetrees, and Divergence Times. Molecular Biology and Evolution 34, 1812–1819, doi:10.1093/molbev/msx116 (2017).

26 Schaum, N. et al. Single-cell transcriptomics of 20 mouse organs creates a Tabula Muris. Nature 562, 367–372, doi:10.1038/s41586-018-0590-4 (2018).

27 Jean-Baptiste, K. et al. Dynamics of gene expression in single root cells of Arabidopsis thaliana. The Plant Cell 31, 993–1011 (2019).

28 Denyer, T. et al. Spatiotemporal developmental trajectories in the Arabidopsis root revealed using high-throughput single-cell RNA sequencing. Developmental cell 48, 840–852. e845 (2019).

29 Ryu, K. H., Huang, L., Kang, H. M. & Schiefelbein, J. Single-cell RNA sequencing resolves molecular relationships among individual plant cells. Plant physiology 179, 1444–1456 (2019).

30 Shulse, C. N. et al. High-throughput single-cell transcriptome profiling of plant cell types. Cell reports 27, 2241–2247. e2244 (2019).

31 Ashburner, M. et al. Gene ontology: tool for the unification of biology. The Gene Ontology Consortium. Nat Genet 25, 25–29, doi:10.1038/75556 (2000).

32 Nehrt, N. L., Clark, W. T., Radivojac, P. & Hahn, M. W. Testing the Ortholog Conjecture with Comparative Functional Genomic Data from Mammals. PLOS Computational Biology 7, e1002073, doi:10.1371/journal.pcbi.1002073 (2011).

33 Adams, K. L. & Wendel, J. F. Polyploidy and genome evolution in plants. Current opinion in plant biology 8, 135–141 (2005).

34 Ohno, S. Evolution by gene duplication. (Springer-Verlag, 1970).

35 Consortium, G. T. The Genotype-Tissue Expression (GTEx) project. Nat Genet 45, 580–585, doi:10.1038/ng.2653 (2013).

36 Liebeskind, B. J., McWhite, C. D. & Marcotte, E. M. Towards Consensus Gene Ages. Genome Biology and Evolution 8, 1812–1823, doi:10.1093/gbe/evw113 (2016).

37 Mustafin, Z. S. et al. Phylostratigraphic Analysis Shows the Earliest Origination of the Abiotic Stress Associated Genes in A. thaliana. Genes 10, doi:10.3390/genes10120963 (2019).

38 Drost, H.-G., Gabel, A., Grosse, I. & Quint, M. Evidence for Active Maintenance of Phylotranscriptomic Hourglass Patterns in Animal and Plant Embryogenesis. Molecular Biology and Evolution 32, 1221–1231, doi:10.1093/molbev/msv012 (2015).

39 Quint, M. et al. A transcriptomic hourglass in plant embryogenesis. Nature 490, 98–101, doi:10.1038/nature11394 (2012).

40 Arendsee, Z. W., Li, L. & Wurtele, E. S. Coming of age: orphan genes in plants. Trends in plant science 19, 698–708, doi:10.1016/j.tplants.2014.07.003 (2014).

41 Hart, T., Brown, K. R., Sircoulomb, F., Rottapel, R. & Moffat, J. Measuring error rates in genomic perturbation screens: gold standards for human functional genomics. Molecular systems biology 10, 733, doi:10.15252/msb.20145216 (2014).

42 Cherry, J. M. et al. Saccharomyces Genome Database: the genomics resource of budding yeast. Nucleic acids research 40, D700–705, doi:10.1093/nar/gkr1029 (2012).

43 Deutekom, E. S., Snel, B. & van Dam, T. J. P. Benchmarking orthology methods using phylogenetic patterns defined at the base of Eukaryotes. Briefings in bioinformatics, doi:10.1093/bib/bbaa206 (2020).

44 Hu, Y. et al. An integrative approach to ortholog prediction for disease-focused and other functional studies. BMC bioinformatics 12, 357–357, doi:10.1186/1471-2105-12-357 (2011).

45 Glover, N. et al. Advances and Applications in the Quest for Orthologs. Molecular Biology and Evolution 36, 2157–2164, doi:10.1093/molbev/msz150 (2019).

46 Altenhoff, A. M. et al. Standardized benchmarking in the quest for orthologs. Nature methods 13, 425–430 (2016).

47 Bruford, E. A. et al. The HGNC Database in 2008: a resource for the human genome. Nucleic acids research 36, D445–D448, doi:10.1093/nar/gkm881 (2007).

48 DeLuca, T. F., Cui, J., Jung, J.-Y., St Gabriel, K. C. & Wall, D. P. Roundup 2.0: enabling comparative genomics for over 1800 genomes. Bioinformatics (Oxford, England) 28, 715–716, doi:10.1093/bioinformatics/bts006 (2012).

49 Schreiber, F. & Sonnhammer, E. L. L. Hieranoid: hierarchical orthology inference. Journal of molecular biology 425, 2072–2081, doi:10.1016/j.jmb.2013.02.018 (2013).

50 Huerta-Cepas, J., Capella-Gutiérrez, S., Pryszcz, L. P., Marcet-Houben, M. & Gabaldón, T. PhylomeDB v4: zooming into the plurality of evolutionary histories of a genome. Nucleic acids research 42, D897–D902, doi:10.1093/nar/gkt1177 (2013).

51 Sonnhammer, E. L. & Östlund, G. InParanoid 8: orthology analysis between 273 proteomes, mostly eukaryotic. Nucleic acids research 43, D234–239, doi:10.1093/nar/gku1203 (2015).

52 Altenhoff, A. M. et al. The OMA orthology database in 2018: retrieving evolutionary relationships among all domains of life through richer web and programmatic interfaces. Nucleic acids research 46, D477–D485, doi:10.1093/nar/gkx1019 (2017).

53 Ruzicka, L. et al. The Zebrafish Information Network: new support for non-coding genes, richer Gene Ontology annotations and the Alliance of Genome Resources. Nucleic acids research 47, D867–D873, doi:10.1093/nar/gky1090 (2018).

54 Nevers, Y. et al. OrthoInspector 3.0: open portal for comparative genomics. Nucleic acids research 47, D411–d418, doi:10.1093/nar/gky1068 (2019).

55 Emms, D. M. & Kelly, S. OrthoFinder: phylogenetic orthology inference for comparative genomics. Genome biology 20, 238, doi:10.1186/s13059-019-1832-y (2019).

56 Yates, A. D. et al. Ensembl 2020. Nucleic acids research 48, D682–D688, doi:10.1093/nar/gkz966 (2019).

57 Mi, H., Muruganujan, A., Ebert, D., Huang, X. & Thomas, P. D. PANTHER version 14: more genomes, a new PANTHER GO-slim and improvements in enrichment analysis tools. Nucleic acids research 47, D419–D426, doi:10.1093/nar/gky1038 (2018).

58 Ruan, J. et al. TreeFam: 2008 Update. Nucleic acids research 36, D735–740, doi:10.1093/nar/gkm1005 (2008).

59 Hulsen, T., Huynen, M. A., de Vlieg, J. & Groenen, P. M. A. Benchmarking ortholog identification methods using functional genomics data. Genome biology 7, R31, doi:10.1186/gb-2006-7-4-r31 (2006).

60 Kachroo, A. H. et al. Systematic humanization of yeast genes reveals conserved functions and genetic modularity. Science (New York, N.Y.) 348, 921–925, doi:10.1126/science.aaa0769 (2015).

61 Laurent, J. M. et al. Humanization of yeast genes with multiple human orthologs reveals functional divergence between paralogs. PLoS Biology 18, e3000627 (2020).

62 Chorostecki, U., Molina, M., Pryszcz, L. P. & Gabaldón, T. MetaPhOrs 2.0: integrative, phylogeny-based inference of orthology and paralogy across the tree of life. Nucleic acids research 48, W553–W557, doi:10.1093/nar/gkaa282 (2020).

63 Crow, M., Paul, A., Ballouz, S., Huang, Z. J. & Gillis, J. Characterizing the replicability of cell types defined by single cell RNA-sequencing data using MetaNeighbor. Nature communications 9, 884, doi:10.1038/s41467-018-03282-0 (2018).

64 Crow, M. & Gillis, J. Co-expression in Single-Cell Analysis: Saving Grace or Original Sin? Trends in genetics : TIG 34, 823–831, doi:10.1016/j.tig.2018.07.007 (2018).

65 Heimberg, G., Bhatnagar, R., El-Samad, H. & Thomson, M. Low Dimensionality in Gene Expression Data Enables the Accurate Extraction of Transcriptional Programs from Shallow Sequencing. Cell Syst 2, 239–250, doi:10.1016/j.cels.2016.04.001 (2016).

66 Koepfli, K. P., Paten, B. & O’Brien, S. J. The Genome 10K Project: a way forward. Annual review of animal biosciences 3, 57–111, doi:10.1146/annurev-animal-090414-014900 (2015).

67 Guan, Y. et al. Tissue-Specific Functional Networks for Prioritizing Phenotype and Disease Genes. PLOS Computational Biology 8, e1002694, doi:10.1371/journal.pcbi.1002694 (2012).

68 Crow, M., Paul, A., Ballouz, S., Huang, Z. J. & Gillis, J. Exploiting single-cell expression to characterize co-expression replicability. Genome biology 17, 101, doi:10.1186/s13059-016-0964-6 (2016).

69 Tarashansky, A. J. et al. Mapping single-cell atlases throughout Metazoa unravels cell type evolution. bioRxiv, 2020.2009.2028.317784, doi:10.1101/2020.09.28.317784 (2020).

70 Vallejo, A. F. et al. Resolving cellular systems by ultra-sensitive and economical single-cell transcriptome filtering. bioRxiv, 800631, doi:10.1101/800631 (2019).

71 Pereira, C., Denise, A. & Lespinet, O. A meta-approach for improving the prediction and the functional annotation of ortholog groups. BMC genomics 15 Suppl 6, S16–S16, doi:10.1186/1471-2164-15-S6-S16 (2014).

72 Ballouz, S., Weber, M., Pavlidis, P. & Gillis, J. EGAD: ultra-fast functional analysis of gene networks. Bioinformatics (Oxford, England) 33, 612–614, doi:10.1093/bioinformatics/btw695 (2017).

73 Ballouz, S. & Gillis, J. Strength of functional signature correlates with effect size in autism. Genome Med 9, 64, doi:10.1186/s13073-017-0455-8 (2017).

74 Yanai, I. et al. Genome-wide midrange transcription profiles reveal expression level relationships in human tissue specification. Bioinformatics (Oxford, England) 21, 650–659, doi:10.1093/bioinformatics/bti042 (2005).

75 McCarthy, D. J., Campbell, K. R., Lun, A. T. L. & Wills, Q. F. Scater: pre-processing, quality control, normalization and visualization of single-cell RNA-seq data in R. Bioinformatics (Oxford, England) 33, 1179–1186, doi:10.1093/bioinformatics/btw777 (2017).

76 Smyth, G. K. in Bioinformatics and computational biology solutions using R and Bioconductor 397–420 (Springer, 2005).

77 Edgar, R., Domrachev, M. & Lash, A. E. Gene Expression Omnibus: NCBI gene expression and hybridization array data repository. Nucleic acids research 30, 207–210, doi:10.1093/nar/30.1.207 (2002).

78 Haghverdi, L., Lun, A. T. L., Morgan, M. D. & Marioni, J. C. Batch effects in single-cell RNA-sequencing data are corrected by matching mutual nearest neighbors. Nature biotechnology 36, 421–427, doi:10.1038/nbt.4091 (2018).

79 umap: Uniform Manifold Approximation and Projection v. R package version 0.2.6.0 (2020).

80 Durinck, S. et al. BioMart and Bioconductor: a powerful link between biological databases and microarray data analysis. Bioinformatics (Oxford, England) 21, 3439–3440 (2005).

