## Supplementary Material for "Coexpression reveals conserved mechanisms of transcriptional cell identity"

### Supplementary Figure 1 – Related to Figure 2

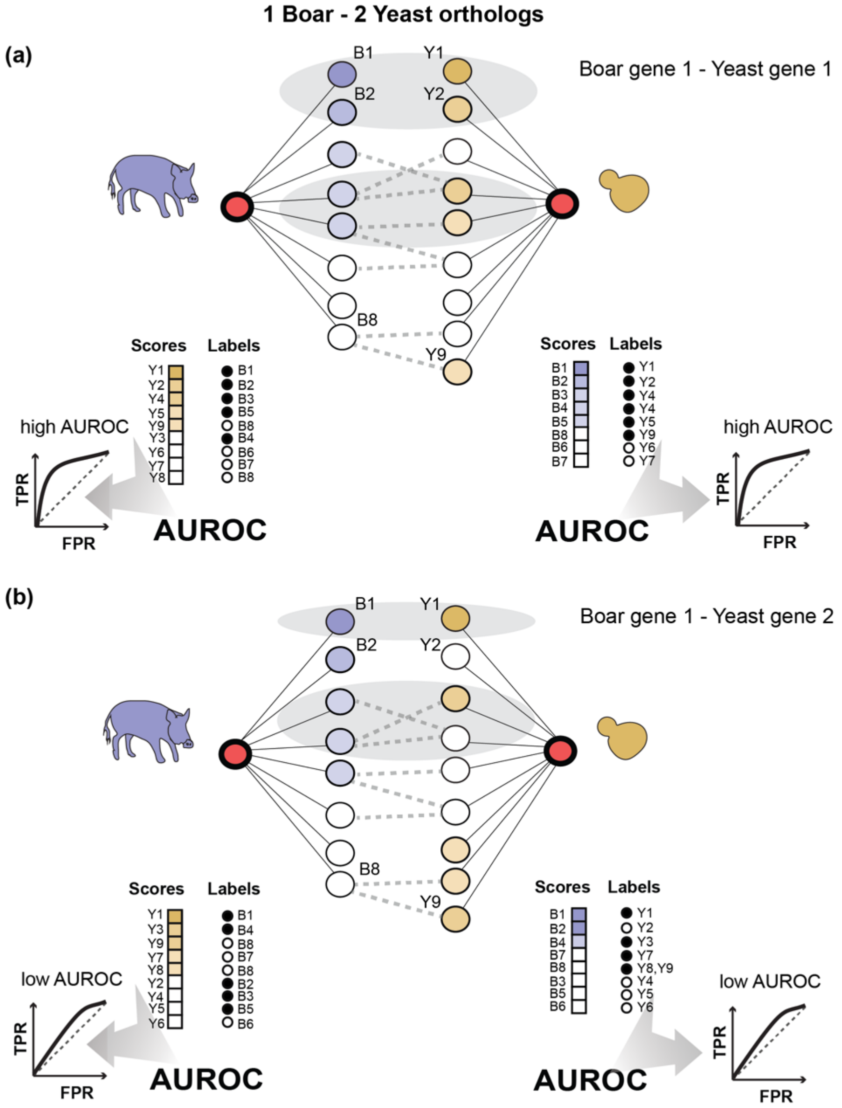

***Schematic of the M-to-N coexpression conservation method***

*This figure illustrates the many-to-many scoring scheme using the example of an orthologous gene group containing 1 boar and 2 yeast orthologs. In the top panel (a) we consider the boar gene, and the first of the two yeast genes, and in the bottom panel (b) we consider the boar gene and the second of the yeast genes. The genes in the orthogroup are colored in red, and the coexpression neighborhoods of those genes are indicated in blue and yellow for boar and yeast respectively. In (a), we see 5 genes in the boar's coexpression neighborhood. These map via orthology relationships to 6 yeast genes, shown with dashed lines when there is not a 1-1 pairing. We see that many of the yeast genes are also strongly coexpressed with the yeast gene in the boar-yeast orthogroup, yielding a high coexpression conservation score (AUROC). In the bottom panel, we illustrate a different scenario. Here, the yeast ortholog's coexpression neighborhood does not align with that of the boar, yielding a low AUROC.*

### Supplementary Figure 2 – Related to Figure 3

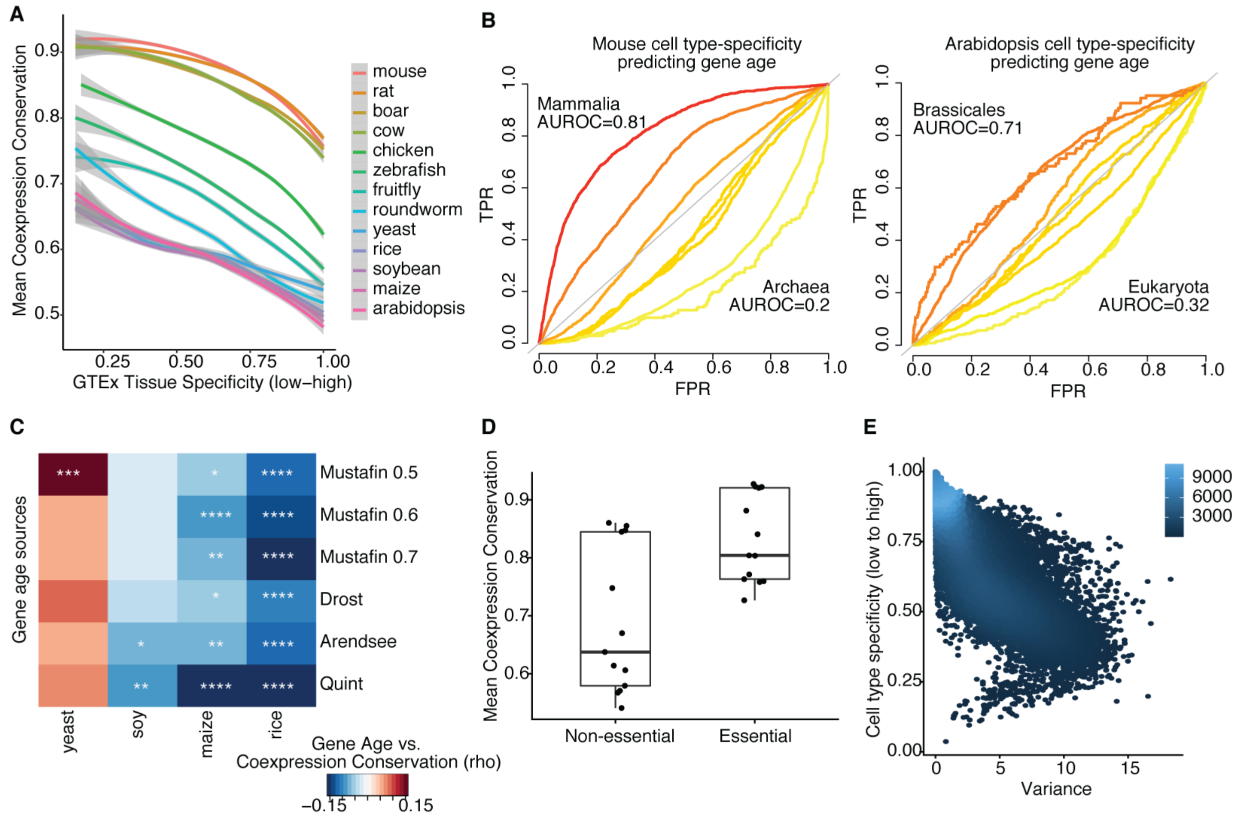

***Gene features associate with coexpression conservation***

**A –** *Coexpression conservation is plotted with respect to human tissue specificity. Lines are loess fits on mean values for each species, +/- SD. Tissue specificity is negatively associated with coexpression conservation.* **B –** *Lines indicate predictions for genes of different ages using mouse (left) or Arabidopsis (right) cell type specificity. Colors represent AUROCs. Younger genes are well predicted by cell type specificity, whereas older genes are negatively predicted.* **C –** *Heatmap of Spearman correlation coefficients between gene age predictions from 6 sources (y-axis) and Arabidopsis coexpression conservation for each of 4 species (1-to-1 orthologs only). Stars indicate p-values (*<0.05, **<0.01, ***<0.001, ****<0.0001). Coexpression conservation is negatively correlated with gene age for the three plants, but not yeast. Some variation is observed across gene age sources.* **D –** *Points show average coexpression conservation between humans and each of the other species, split by whether the genes are known to be essential in human or not. Essential genes have significantly higher coexpression conservation (p<0.01).* **E –** *Scatterplot shows genes plotted with respect to their cell type specificity and pseudo-bulk variance in Tabula Muris. Colors indicate the local point density. Cell type specificity is negatively associated with variance.*

### Supplementary Table 1 – Related to Figure 4

For each orthology prediction algorithm with data for all species in the Alliance of Genome Resources database, mean and standard deviation of coexpression conservation scores are listed. OrthoDB is also included. Algorithms appear in descending order based on mean coexpression conservation.

| Orthology Algorithm | Mean Coexpression Conservation | Standard Deviation |
|---|---|---|
| **Hieranoid** | 0.75 | 0.16 |
| **InParanoid** | 0.75 | 0.16 |
| **OMA** | 0.74 | 0.16 |
| **OrthoInspector** | 0.74 | 0.16 |
| **TreeFam** | 0.74 | 0.16 |
| **OrthoDB** | 0.73 | 0.13 |
| **OrthoFinder** | 0.73 | 0.16 |
| **PhylomeDB** | 0.73 | 0.16 |
| **Ensembl Compara** | 0.72 | 0.16 |
| **PANTHER** | 0.72 | 0.16 |

### Supplementary Table 2 – Related to Figure 4

This table provides coexpression conservation scores for yeast-human gene pairs tested for complementarity in Kachroo et al. Data can be found in a separately uploaded Excel spreadsheet. Column descriptions are in the first worksheet, and data in the second worksheet. Gene pairs appear in order of descending coexpression conservation.

### Supplementary Note – CoCoCoNet Tutorial

##### *Purpose and scope*

The purpose of CoCoCoNet is to provide access to gold-standard coexpression networks across a diverse set of species, enabling comparative coexpression analyses.

##### *Tutorial*

CoCoCoNet requires inputting a list of genes, or a single gene, as gene symbols, Ensembl ID, or NCBI Entrez ID, and the corresponding species to be used in the construction of the network. After this, users can select optional parameters before visualizing and downloading results.

##### *Initialization*

Preloaded genes can be used to test the server's functionality. In this tutorial we will use a random sample of 250 essential mouse genes curated by the MacArthur lab from the Mouse Genome Database (https://github.com/macarthur-lab/gene_lists) by clicking the appropriate box under "Example settings". In the boxes on the right, users can input genes by typing them, selecting them from a drop-down list or by uploading a text file with genes separated by lines.

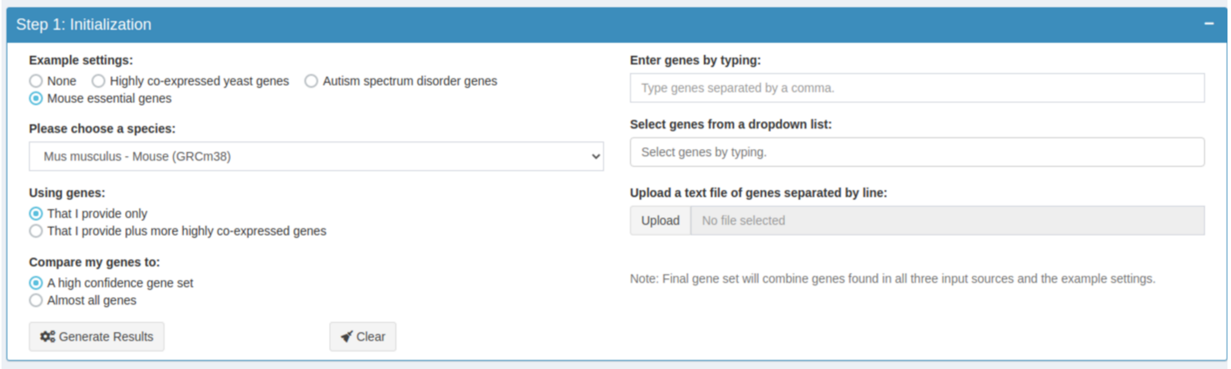

***Figure 1:*** *Screenshot of the CoCoCoNet interface. Selected options are used in this tutorial.*

Analyses can be extended beyond the input gene set with the "Using genes:" options.

- Selecting "That I provide only" will construct a network using only selected genes.
- Selecting "That I provide plus more highly co-expressed genes" will allow you to choose how many more of the most highly correlated genes to use that are not in the provided set.

The "Compare my genes to:" options allow the user to limit the genes to a high confidence set or all genes based on expression levels.

- Selecting “A high confidence gene set” will match input genes to only a subset of genes filtered on minimum expression level across experiments.
- Selecting “Almost all genes” will match your genes to a very lightly filtered gene set.

##### *Generating Results*

Once a set of genes has been input, selecting “Generate Results” will display a coexpression network (**Figures 2 and 3**), and the distribution of coexpression values. A sliding threshold bar is provided to control the sparsity of the visualization, allowing the user to focus on the most co-expressed gene pair relationships. Selecting a node on the network will display a table of the selected gene’s top co-expressed pairs, their values and a short summary of the gene if available (**Figure 2**).

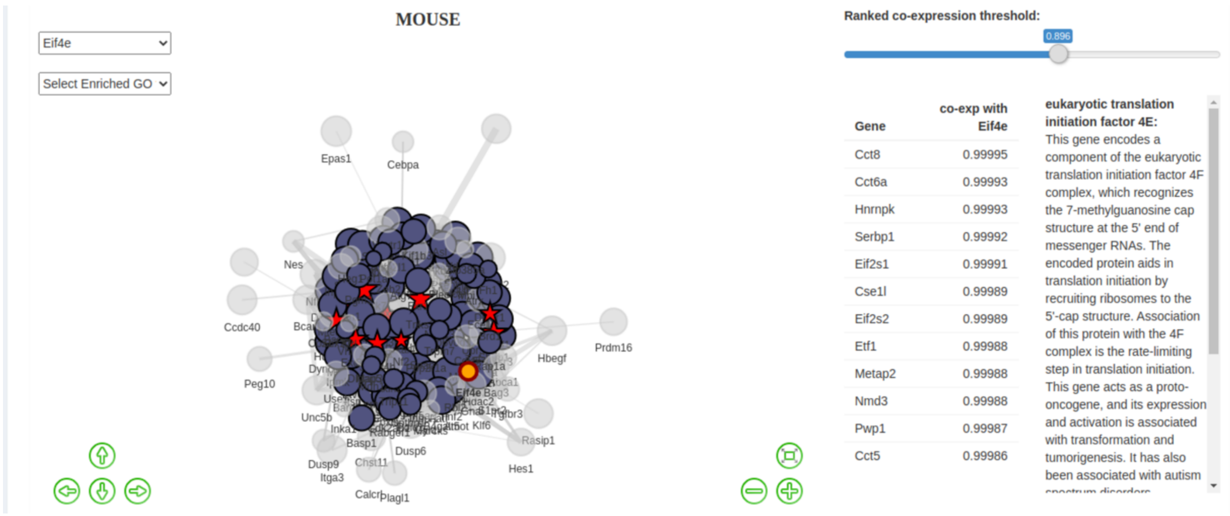

***Figure 2:*** *Generated results using a random sample of 250 essential mouse genes. Selecting a gene on the network will display the top co-expressed genes, and a brief summary of the gene if it’s available.*

##### *Conserved species comparison*

The next section allows the user to select a second species to compare to the first. Choosing a species and generating results will search for orthologous genes from OrthoDB and display the coexpression network from those genes. The user is then able to select genes to view additional information, highlight genes by GO annotation, or, toggle between using all orthologs or restricting to 1-to-1 orthologs. Next, the distributions of coexpression conservation scores are displayed as a histogram (**Figure 4**). A data table and visualization of specific genes are available, allowing in-depth exploration of coexpression conservation. Users can select a gene to display its coexpression conservation score, along with the scores of its top coexpressed gene pairs.

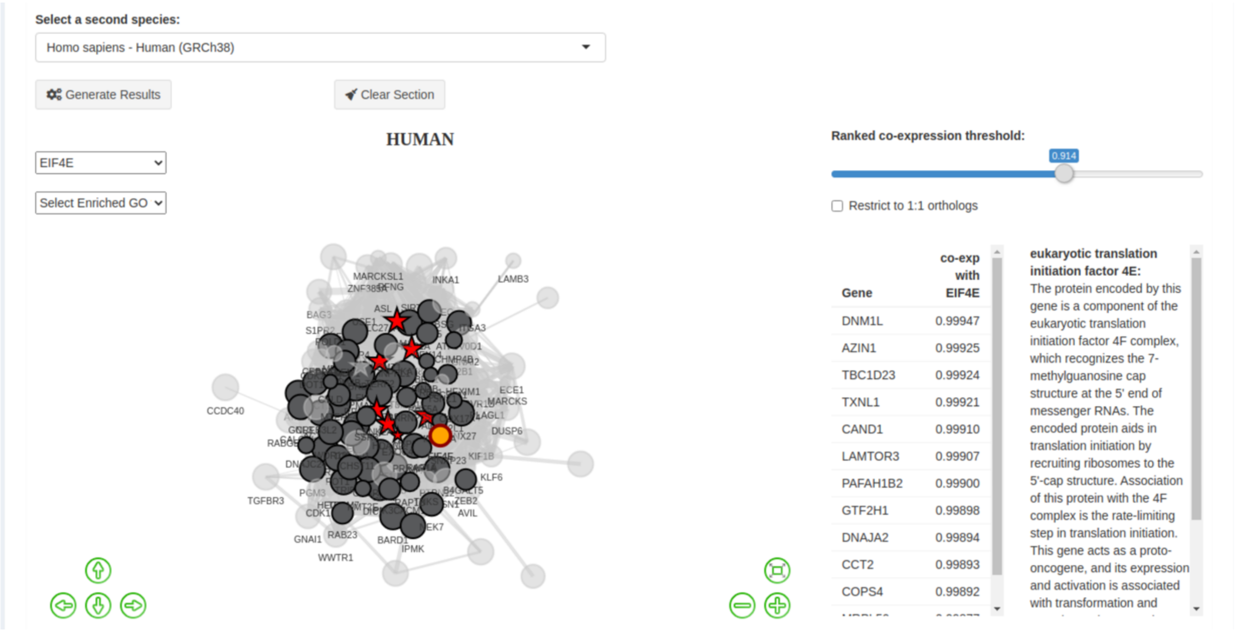

***Figure 3****: Human genes were selected to compare to the set of essential mouse genes.*

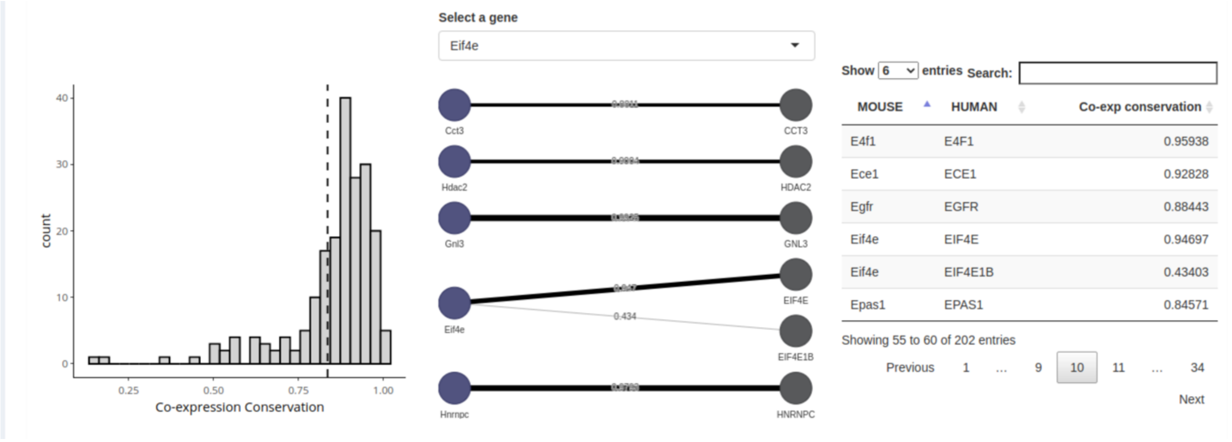

***Figure 4:*** *(Left) Distribution of coexpression conservation scores. (Middle) A visualization of Eif4e and its top coeexpressed genes in mouse along with the coexpression conservation scores for each in human. (Right) Scores are also available as a table that can be sorted.*

##### *Exporting Results*

Results can be downloaded at each step by selecting the ‘Download’ option. Users are also able to download all of the relevant data, including their gene list, gene pairs within the network, GO enrichment scores, and GO annotations, ortholog mappings, coexpression conservation, TAGR annotations, and more. Networks can be exported by right-clicking and saving the image, and other figures can be exported by selecting the plotly download button (**Figure 5**). Data can also be downloaded in bulk from ftp://milton.cshl.edu/data.

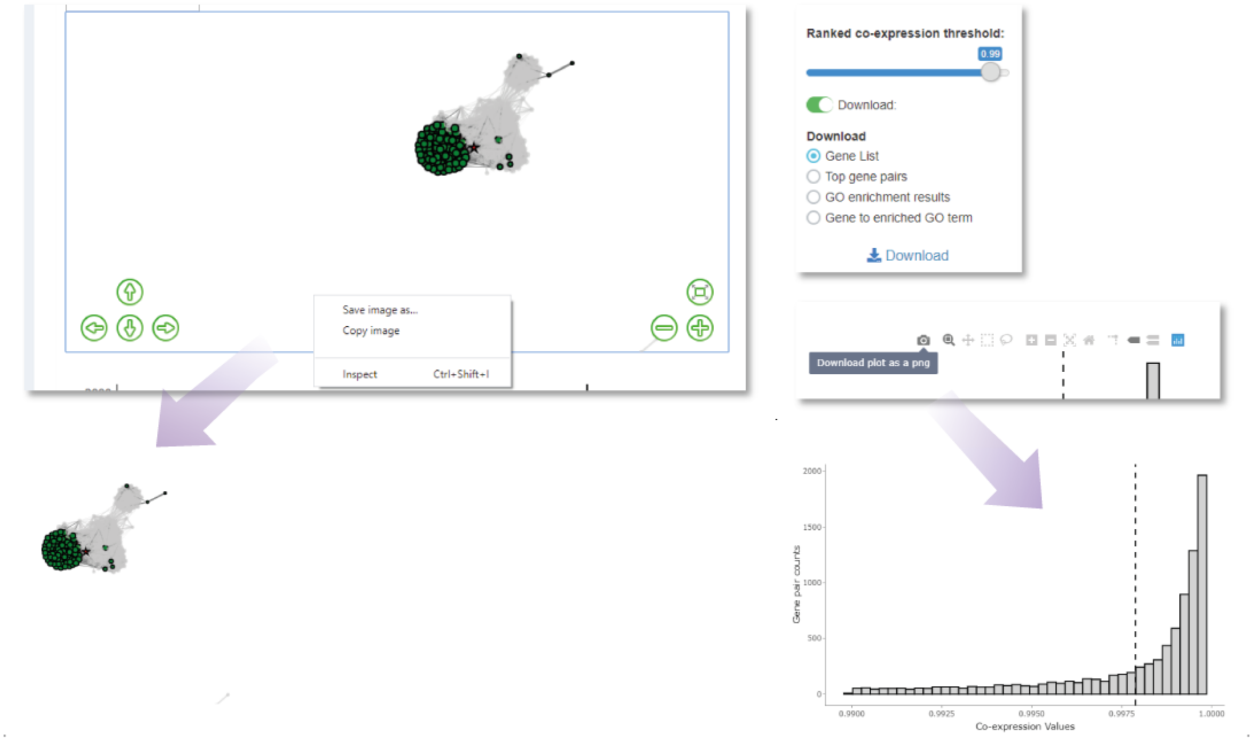

***Figure 5:*** *Data and figures are easily downloadable at each stage of the analysis.*
